# Expression of cell-wall related genes is highly variable and correlates with sepal morphology

**DOI:** 10.1101/2022.04.26.489498

**Authors:** Diego A. Hartasánchez, Annamaria Kiss, Virginie Battu, Charline Soraru, Abigail Delgado-Vaquera, Florian Massinon, Marina Brasó-Vives, Corentin Mollier, Marie-Laure Martin-Magniette, Arezki Boudaoud, Françoise Monéger

## Abstract

Control of organ morphology is a fundamental feature of living organisms. There is, however, observable variation in organ size and shape within a given genotype. Taking the sepal of Arabidopsis as a model, we investigated whether we can use variability of gene expression alongside variability of organ morphology to identify gene regulatory networks potentially involved in organ size and shape determination. We produced a dataset composed of morphological parameters and genome-wide transcriptome obtained from 27 individual sepals from wild-type plants with nearly identical genetic backgrounds, environment, and developmental stage. Sepals exhibited appreciable variability in both morphology and transcriptome, with response to stimulus genes and cell-wall related genes displaying high variability in expression. We additionally identified five modules of co-expressed genes which correlated significantly with morphology, revealing biologically relevant gene regulatory networks. Interestingly, cell-wall related genes were overrepresented in two of the top three modules. Overall, our work highlights the benefit of using coupled variation in gene expression and phenotype in wild-type plants to shed light on the mechanisms underlying organ size and shape determination. Although causality between gene expression and sepal morphology has not been established, our approach opens the way to informed analysis for mutant characterization and functional studies.

## Introduction

Size and shape of organs are very important traits that determine adaptation to a given environment. Accordingly, numerous studies have addressed the developmental regulation of size and shape. For instance, screens for mutants with altered organ size have allowed the identification of mutants with defects in cell size, cell number, or both (reviewed in Czesnick & Lenhard, 2015). The characterisation of such mutants has highlighted the contribution of individual genes to the regulation of size or shape. However, genes are integrated in complex molecular networks that operate in each cell and drive cellular behavior. As a result, events occurring at the molecular level control phenomena occurring at a higher scale (Long and Boudaoud, 2019). This contributes to the difficulty of deciphering the mechanisms underlying morphogenesis. Despite enormous progress in understanding how gene expression patterns are established during development (Alvarez-Buylla, et al., 2008; Liu et al., 2020), we still need to understand how these patterns are related to cellular processes leading to the development of organs with robust size and shape (e.g. Zhu et al., 2020). In this context, modules of interacting genes appear as a central layer of biological organization (Lucas et al., 2011), calling for new approaches to identify such modules.

The association between genotype and phenotype can be assessed in many different ways. A routinely used approach is based on the characterization of the effects of a mutation on the phenotype of the organism. However, its main drawback is that, because genes are part of complex regulatory networks, it may be hard to draw reliable information from the phenotype of one- or two-gene mutants, especially when studying complex traits (Chen et al., 2018). In addition, mutations often induce compensation that increases the difficulty to understand the link between genotype and phenotype (reviewed in Hisanaga et al., 2015). Regarding the choice of scale, single-cell gene expression data has enabled the characterization of complex organs with multiple different cell-types such as the root (Dorrity et al., 2021) and ovules (Hou et al., 2021) in Arabidopsis. Complementarily, transcriptomic data at the organismal level, such as that obtained from whole seedlings, has provided very valuable insight regarding variability in gene expression (Cortijo et al., 2019) highlighting the role of modules of co-expressed genes (Cortijo et al., 2020). Both of these approaches, however, have their shortcomings. On the one hand, single-cell RNA sequencing has limited power to detect lowly expressed genes. On the other hand, transcriptomes obtained from whole individuals combine different structures, tissues and cell-types. Here, we have chosen an intermediate scale by studying a whole organ, the sepal, which has relatively few cell-types.

Recently, the Arabidopsis sepal has emerged as a good system to study organ morphology (Roeder, 2021) for several reasons: Arabidopsis has a large number of flowers on each inflorescence; sepals are very accessible for dissection, imaging and experimentation; and sepals exhibit very reproducible morphology. Here, we use the sepal of *Arabidopsis thaliana* to explore wild-type variability in morphology and in gene expression. We use this variability to identify modules of co-expressed genes to reconstitute gene regulatory networks (GRNs) associated with sepal morphology. We have performed high-coverage RNA sequencing to obtain high-quality transcriptomic data at the organ level, aiming at identifying modules of co-expressed genes which could be linked to sepal morphology. To identify these modules in wild-type plants, we have evaluated variation in sepals by obtaining gene expression data and morphology from 27 wild-type sepals in environmentally controlled conditions and with identical genetic background. Taking advantage of molecular and phenotypic variation among a high number of wild-type sepals, we extracted relevant biological information about cell-wall related genes and organ morphology.

## Results

### A unique dataset enables the analysis of gene expression and morphology of single sepals

Our goal was to analyze the variability of both the morphology and the transcriptome of individual sepals from wild-type plants grown in standard conditions. On a single inflorescence (flowering stem) of Arabidopsis, flower buds initiate at regular time intervals in a continuous manner. Each flower bud contains four sepals that are formed in a defined order, the first one to emerge being called the abaxial sepal (it forms on the side furthest away from the flower meristem and is also known as the outer sepal). Previous studies (Hervieux et al., 2016; Hong et al., 2016) have shown that arrest of sepal growth begins at stage 11 (according to staging of Arabidopsis flower development described in Smyth et al., 1990). We therefore chose early stage 11 to collect sepals and to minimize stage heterogeneity. Indeed, early stage 11 is rather transient, as we rarely find two flowers at this stage in a given inflorescence. We then collected 30 individual abaxial sepals, each from 30 different secondary inflorescences, taken from three different Col-0 wild-type plants (10 sepals from each plant), labeled D, E and F, grown simultaneously in the same standard experimental conditions, and used them to recover their 3D shape as well as their RNA to perform RNA-Seq analysis (Figure 1A). Each sepal was imaged under a confocal microscope using autofluorescence before being frozen in liquid nitrogen for RNA extraction, less than 5 minutes after the dissection. Sepal images were analyzed with in-house python scripts and using MorphographX (Barbier de Reuille et al., 2015; see Materials and Methods). Our image analysis pipeline allows for automated, precise and quantitative analysis of sepal shape compatible with large-scale analysis, which can be used to characterize new mutants as well as to re-examine known mutants. One advantage of the protocol is that it exploits the natural autofluorescence of plant tissue and does not require introducing any fluorescent protein by genetic transformation, like in previous work on leaves (Biot et al., 2016).

**Figure 1:**
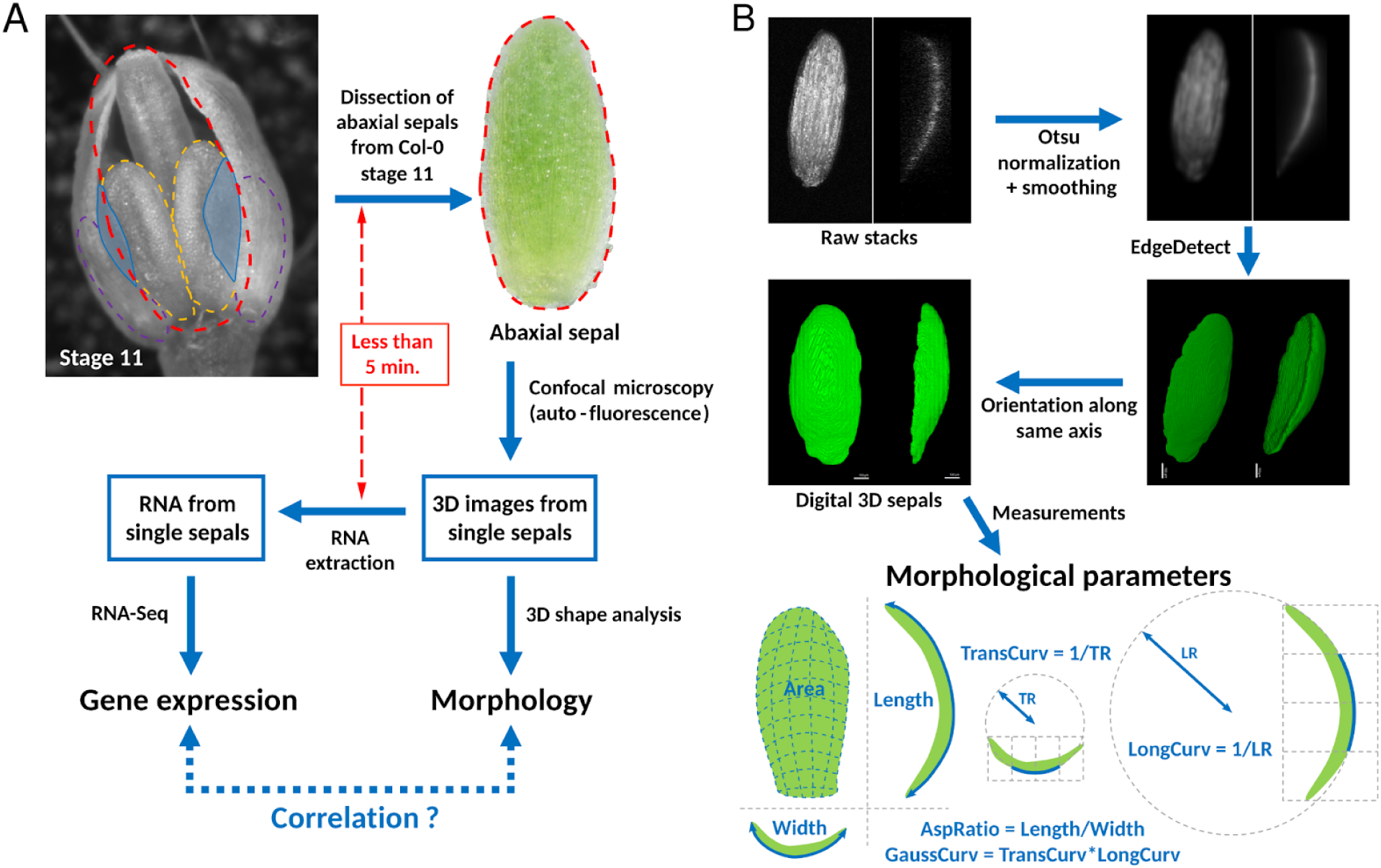
Protocol used for sample collection, data generation and image analysis. (A) Flower buds at early stage 11 were identified as such when petals (in blue) were longer than the lateral stamen (outlined by purple dashed lines, not visible on the image) but shorter than the median stamen (yellow dashed lines). The abaxial sepal was dissected and put on agar to avoid dehydration. Autofluorescence enabled imaging of the entire sepal using a confocal microscope with the lowest possible laser power and resolution to minimize laser exposure. Within less than 5 minutes, the sepal was frozen in liquid nitrogen. After grinding, RNA was extracted from individual sepals and used for RNA-Seq. (B) In order to retrieve precise morphological parameters for each sepal sample, raw images obtained from the confocal microscope were normalized, smoothed, and segmented into 3D volumes representing sepals, before fixing sepal orientation. From the digital 3D images, morphological parameters were extracted, namely, length, width, aspect ratio, area, and transversal, longitudinal and Gaussian curvatures (see Materials and Methods). Transversal (longitudinal) curvature was measured from the central zone of the sepal by dividing the total width (length) of the sepal in four equal sections as shown in the figure, then, by fitting the circumference that best matched the sepal curvature in the middle two sections and taking the inverse of the radius of said circumference.

Sepal morphology was characterized by measuring curvilinear length (Length), curvilinear width (Width), surface area (Area), longitudinal curvature (LongCurv) and transversal curvature (TransCurv) (Figure 1B). From these measurements, we additionally calculated the aspect ratio (AspRatio) by dividing Length over Width, and the Gaussian curvature (GaussCurv) by multiplying LongCurv by TransCurv. The parameters that we have used here to describe sepal morphology were selected after considering several parameters and carefully evaluating their reproducibility, in particular by measuring the same sepal several times (see Materials and Methods). Other parameters that had originally been measured, such as curvature at the borders, were excluded because they were not reliable. For this reason, curvature was measured from the central zone of the sepal (Figure 1B). From our transcriptome pre-analysis (see Materials and Methods), we removed lowly expressed genes and restricted the analysis to a set of 14,085 genes with reliable quantification of expression (Supplementary Table 1). We detected three outliers (all from one plant, plant F) which were not further analyzed. All results shown hereafter are for 27 sepal samples.

### Variability in wild-type sepal morphology is mostly explained by variation in area and aspect ratio

We analyzed the seven morphological parameters mentioned above (Length, Width, AspRatio, Area, LongCurv, TransCurv and GaussCurv) within plants and across our 27 sepals (Supplementary Table 2). Analysis of correlations between each pair of parameters (Figure 2A) shows that Area is the most informative parameter since it is strongly positively correlated with Length and Width (as expected, even though it is an independent measurement) and negatively correlated with all three curvature measurements, particularly with GaussCurv. These correlations are statistically significant when considering all 27 samples, and also when evaluating plants independently, despite the expected smaller statistical significance found for plant F due to its smaller sample size (from exclusion of the three outlier sepals according to the RNA-Seq data). In general, anticorrelations observed between size and curvature measurements indicate that larger sepals tend to be less curved in their central region, a result which was beyond reach with previous 2D descriptions of flattened sepals (e.g. Hong et al., 2016).

**Figure 2.**
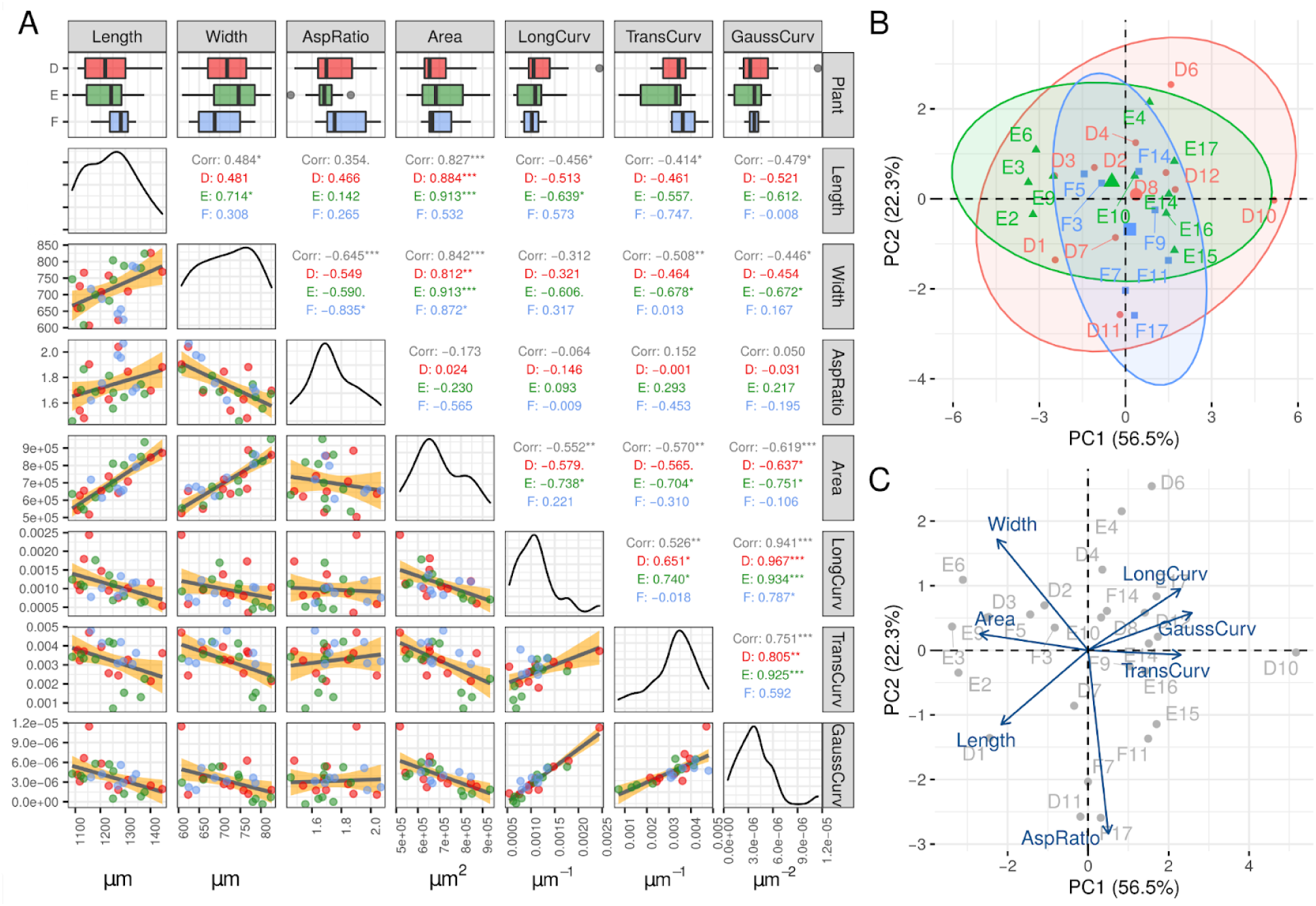
Pairwise comparison and principal component analysis of morphological parameters (A) Comparison between morphological parameters. The first row shows boxplots for each parameter and each plant (D in red, E in green, F in blue), with medians represented with a black line. Colored boxes extend from the first to the third quartile, while whiskers extend a further 1.5 times the interquartile range. Scatter plots for each pairwise comparison show each of the samples with its corresponding color. The gray line with yellow shading corresponds to a linear model adjustment. Pearson correlation coefficients for each pair are shown for all samples pulled together (in gray) and for each plant independently (in green, red and blue) with corresponding p-values shown with asterisks (*** implies p-value < 0.001, ** implies p-value < 0.01, * implies p-value < 0.05, and . implies p-value < 0.1). The diagonal shows the probability density plot for each parameter. (B) Principal component analysis (PCA) for all samples according to their morphological parameters (D in red, E in green, F in blue), showing the percentage of variance explained by the first (PC1) and second (PC2) principal components. Ellipses show 95% confidence ellipses for each set of 2D normally distributed samples for each plant in the corresponding color. (C) Loadings for the PCA shown in B depicting the contribution of each morphological parameter to the first and second principal components in the 2D space.

Performing principal component analysis (PCA) on morphological measurements (Figures 2B & 2C) confirms that despite identical or nearly identical genetic backgrounds, environment and developmental stages between sepal samples, we do observe some level of variability among our samples. The first principal component accounts for 56.5% of the variance and separates Length, Width and Area from LongCurv, TransCurv and GaussCurv, in agreement with correlations observed in Figure 2A. The second principal component, explaining 22.3% of the variance, is mostly dominated by AspRatio, which has no correlation with Area or curvature parameters.

We do not observe any clear clustering by plant although plant F sepals are more distributed along PC2 than PC1 as opposed to plants D and E. This is caused by variability in curvature parameters being reduced and aspect ratio being more variable in plant F sepals compared to the other two plants. From these observations we can conclude that area and aspect ratio are complementary measurements that describe sepal shape and size to a large extent. The third principal component explains 14.1% of the variance and groups all parameters, including Area and curvature parameters, in the same direction with the strongest contribution coming from Length (Supplementary Figure 1).

### Genes with highly variable expression are enriched in response to stimulus and cell-wall functions

Having analyzed the morphological variability present in wild-type sepals, we compared it with variability in gene expression across samples. We calculated the squared coefficient of variation (CV^2^) for each morphological parameter and for each of our 14,085 genes from the RNA-Seq data across 27 samples. CV^2^ (the variance divided by the square of the mean) is a dimensionless quantity that evaluates variability and that allows for comparison between measurements of different nature. Regarding morphological parameters, CV^2^ values for Length, Width and AspRatio are lower than CV^2^ values for curvature parameters, with CV^2^ for Area being between those two groups (Figure 3A). Although there can potentially be intrinsic differences between size and curvature parameters, a possible interpretation of this difference is that the size of the sepal is much more constrained than its curvature. When comparing CV^2^ between morphology and gene expression, as depicted in Figure 3A, we find that Length, Width and AspRatio exhibit less variation (as measured by CV^2^) than gene expression. This observation could be a consequence of noise buffering in gene expression when the latter is translated to phenotype (morphology, in this case).

**Figure 3.**
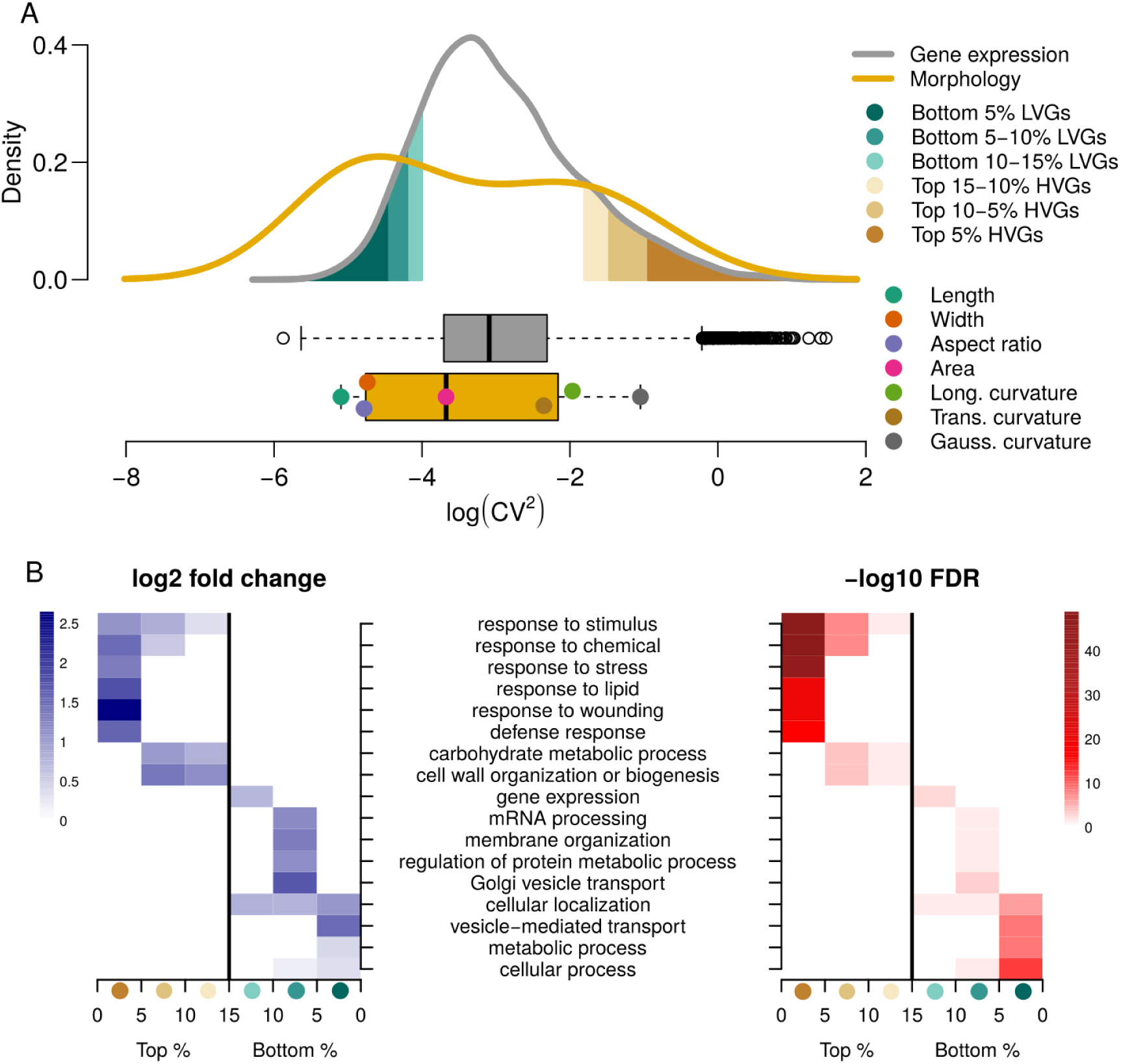
Variability in gene expression and morphology, and GO enrichment of genes grouped by variability in their expression (A) Density profile and boxplots of the squared coefficient of variation (log scale) of gene expression for each of the 14085 genes (gray) and for seven morphological parameters (yellow). Points beyond the whiskers in the upper box plot correspond to outliers. Points over the lower boxplot show the location of CV^2^ values for each morphological parameter. Shades within the gene expression density plot indicate the range of CV^2^ values covered by the selected highly variable and lowly variable gene categories used in B. (B) GO enrichment results for each highly variable and lowly variable gene categories as identified in A. Only top hierarchy GO terms that appear enriched in any of the shown categories are included (see Methods). Log2 fold change enrichment (left) and -log10 false discovery rate (right) are shown for each category.

We then classified genes according to the CV^2^ values of their expression. Given the higher variability (both technical and biological) found for lowly expressed genes, one can correct CV^2^ to account for this association (Cortijo et al., 2019). However, the gene expression threshold that we have set has eliminated most lowly expressed genes from our dataset and essentially, there is no correlation between CV^2^ values and average gene expression (Supplementary Figure 2). Hence, we have ordered genes by the raw CV^2^ values of their expression (across our 27 samples) and have selected highly variable genes (HVGs: top 5%, top 5-to-10%, and top 10-to-15%), and lowly variable genes (LVGs: bottom 5%, bottom 5-to-10%, and bottom 10-to-15%). We verified that the identification of LVGs and HVGs was not exclusively due to differences in gene expression between different plants (Supplementary Figures 3 & 4) by extracting highly and lowly variable genes independently for each plant and comparing the lists obtained between plants and their overlap with the LVG and HVG lists obtained considering all 27 sepal samples. Out of the 704 least variable genes identified in each plant, 89 of them were common when only 2 would be expected if CV^2^ values were randomly assigned to each gene. Reassuringly, all of these 89 genes were among the bottom 5% LVGs identified pooling all 27 sepals together, and only 82 in the bottom 5% LVGs were not identified as lowly variable for any individual plant. Reproducibility for HVGs was much more striking. Out of the 704 most variable genes identified in each plant, 274 of them were common and 272 of them were also in our top 5% HVG set (pooled sepals). Reproducibility of lowly and highly variable genes across plants also holds when taking the top and bottom 15% genes (Supplementary Figures 3 & 4). These results validate the use of CV^2^ across all 27 sepals and the list of genes within the HVG and LVG sets.

In order to see if HVGs and LVGs were enriched in particular biological functions, we searched for enrichment of Gene Ontology (GO) categories in each of these subsets of genes focusing on high hierarchy GO terms using the PANTHER17.0 online tool (Mi et at., 2021; see Materials and Methods). The top 5% HVGs show a very different GO enrichment profile compared to the bottom 5% LVGs (Figure 3B). Top HVGs are enriched in response to stimulus, to chemical, to stress, to lipid, to wounding and defense response, whereas bottom LVGs are enriched in cellular and metabolic processes, vesicle-mediated transport and cellular localization. These findings are consistent with other studies that have found HVGs to be enriched in genes involved in response to environment and LVGs enriched in housekeeping functions (Cortijo et al., 2019; Zheng et al., 2004). An unexpected finding, however, was that within the top 5-to-10% and top 10-to-15% HVGs there is an enrichment in carbohydrate metabolic process and cell-wall organization or biogenesis genes (Figure 3B).

Our results point at HVGs being enriched in cell-wall functions according to GO annotations. However, the latter are known to be incomplete and biased (Timmons et al., 2015). In order to use an alternative approach to assess cell-wall related gene expression in sepals, we compiled a non-exhaustive but rather comprehensive list of cell-wall related genes centered on genes encoding proteins involved in structure, biosynthesis and cell-wall remodeling from the entire Arabidopsis genome (see Materials and Methods). We recovered a list of 1,585 genes (Supplementary Table 3) that we will refer to as our cell-wall related gene (CWRG) list. Among this list, 718 genes are within our 14,085 sepal gene set. Gene expression CV^2^ values of these 718 genes is higher compared to those of the entire gene dataset (Supplementary Figure 5). We confirmed this through a bootstrapping approach by calculating gene expression CV^2^ means for 1000 random samples of 718 genes from the entire dataset to have gene sets of equal size. The gene expression CV^2^ mean of our CWRG list falls clearly above the 95% confidence interval of the distribution corresponding to the entire dataset (Supplementary Figure 5).

### Analysis of co-expression reveals gene modules associated with morphology

We then used the Weighted Correlation Network Analysis (WGCNA) package (Langfelder & Horvath, 2008) to identify modules of co-expressed genes among our 27 samples and evaluated whether some of these modules were significantly associated with morphology (see Materials and Methods). Clustering samples by their gene expression profiles reveals that sepals from the same plant do not have a more similar gene expression profile among them compared to sepals from other plants (Figures 4A & 4B). Tree reconstruction based on gene expression does not yield a clear pattern when normalized morphological parameters are plotted against it (Figure 4B), suggesting a complex link between gene expression and sepal morphology.

**Figure 4:**
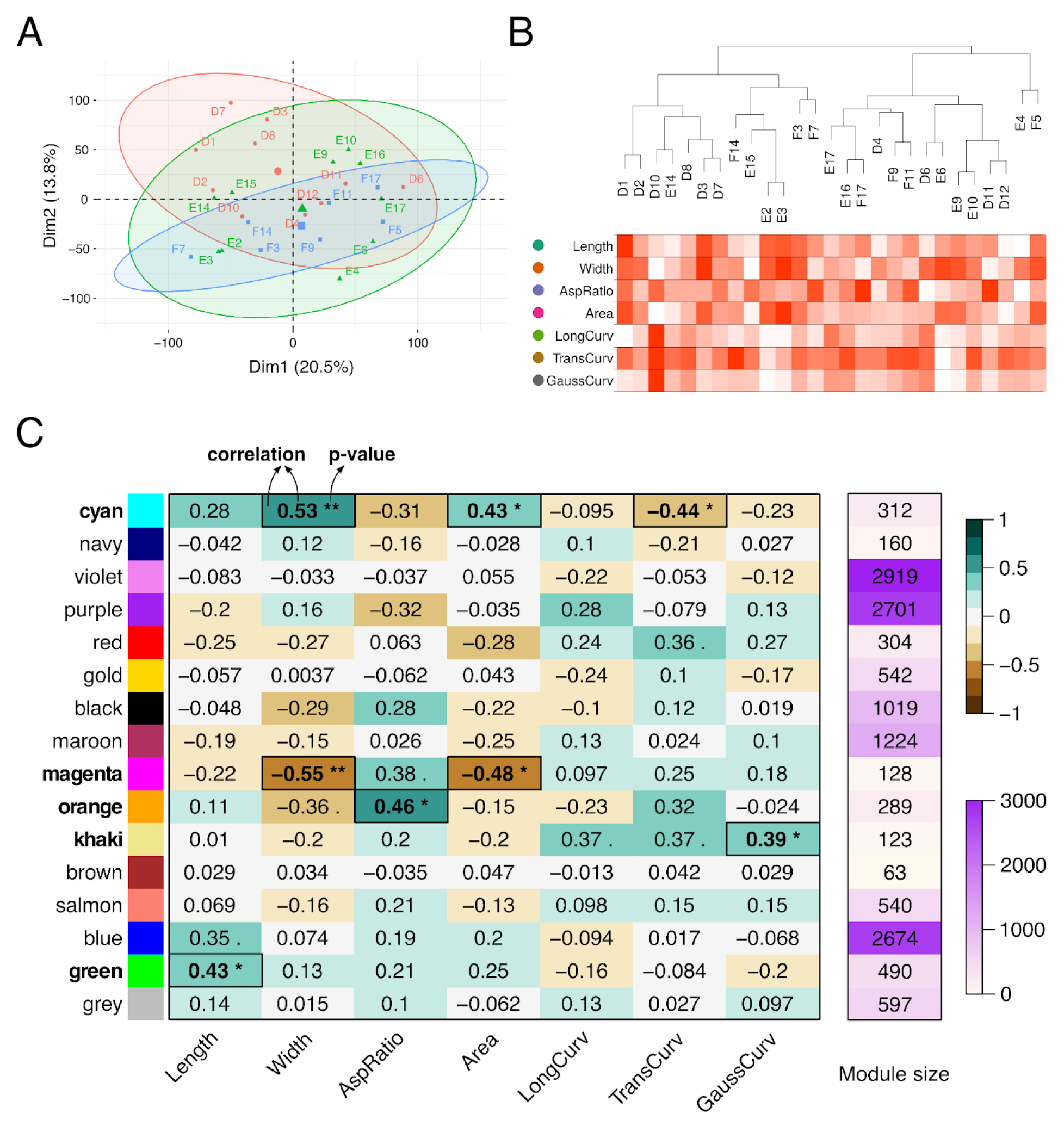
Modules of co-expressed genes associated with morphological parameters (A) PCA on RNA-Seq for 14085 genes across 27 samples belonging to three different plants (D in red, E in green, F in blue) (B) Sample dendrogram from gene expression data reconstructed with WGCNA (Langfelder & Horvath, 2008) and relative magnitude of each morphological parameter measured for each sample (normalized across samples, with intense red depicting the maximum value and white, the minimum value). (C) Module-parameter associations obtained with WGCNA (Langfelder & Horvath, 2008). Modules are randomly assigned a color and are of different sizes, shown on the right panel on a floral white-to-violet color scale. Pearson correlation values between module eigengene expression and morphological parameters (left panel) are shown in black numbers and in a brown-to-green color scale with corresponding p-values shown with asterisks (** implies p-value < 0.01, * implies p-value < 0.05, and . implies p-value < 0.1). Module-parameter pairs with p-values below 0.05 are highlighted with black rectangles with the corresponding module names in bold.

Clustering genes in modules according to their expression patterns across samples and searching for correlations between module eigengene expression (theoretical gene expression profile that is representative of the module) and morphological parameters can allow us to detect gene regulatory networks linked to morphology. We identified 16 modules of co-expressed genes of varying size, ranging from 63 to 2919 genes (Figure 4C, right panel). The list of genes expressed in sepals (14,085 genes) along with the module each gene belongs to is provided in Supplementary Table 4. We found five modules of co-expressed genes whose eigengene correlated with a significant p-value (p < 0.05) with at least one morphological parameter (Figure 4C, left panel). The magenta module has the strongest correlation value (with Width), followed by the cyan module (with Width), the orange module (with AspRatio), the green module (with Length), and the khaki module (with GaussCurv). A visual representation of correlations between modules and morphological parameters is shown in Supplementary Figure 6 by selecting genes with high module membership and high gene significance (in absolute values) for the top-correlated morphological parameter of the corresponding module (see Materials and Methods). Normalized gene expression values for genes with high module membership and high gene significance for 1/Width, Width, and AspRatio in the magenta, cyan and orange modules, respectively, correlate well with the parameter values themselves. Genes with low module membership and gene significance do not correlate as well among each other or with morphological parameters. The green and khaki modules correlate with Length and GaussCurv, respectively, although not as strongly as for the top three modules. These five modules are relatively small in size within the distribution of module sizes (Figure 4C) which is expected for biologically meaningful subsets of genes.

### Modules associated with morphology are enriched in cell-wall related genes

We performed GO enrichment analysis (see Materials and Methods) on all modules (Supplementary Table 5) using the 14,085 gene set as a background, and focusing on the top correlated modules. The magenta module showed an enrichment in cell wall organization or biogenesis and other GO subcategories, such as xylan, cell wall polysaccharide, cell wall macromolecule and hemicellulose metabolic processes, and also secondary cell wall biogenesis, among other terms. The orange module also showed GO enrichment in cell wall organization or biogenesis, but no further GO terms surpassed the FDR < 0.05 threshold with the 14,085 gene set background. To have slightly more power to detect relevant GO terms in the orange module, we performed GO enrichment analysis with the whole genome of Arabidopsis as background (results for the top modules are shown in Supplementary Table 6). Doing so, we found that supramolecular fiber organization, cell morphogenesis, and polysaccharide and pectin biosynthetic processes are enriched in the orange module. The cyan module showed an enrichment peptide biosynthetic process, thylakoid membrane organization and chloroplast rRNA processing, as well as translation. In addition, the green module was enriched in photosynthesis-related GO terms and the khaki module, in response to oxygen levels. We inspected the module membership and gene significance values for the genes associated with the most enriched GO term in each of these five modules (Figure 5). Cell-wall related genes in the magenta module have high module membership and gene significance values suggesting that correlation between the magenta module and Width is driven by cell-wall related genes. The cyan and orange modules also exhibit high module membership for genes associated with peptide biosynthetic process and cell wall organization or biogenesis, respectively, but not particularly high gene significance values. A similar situation can be observed for the green and khaki modules with slightly higher module membership values but average gene significance values of GO-term associated genes with respect to the rest of the module.

**Figure 5:**
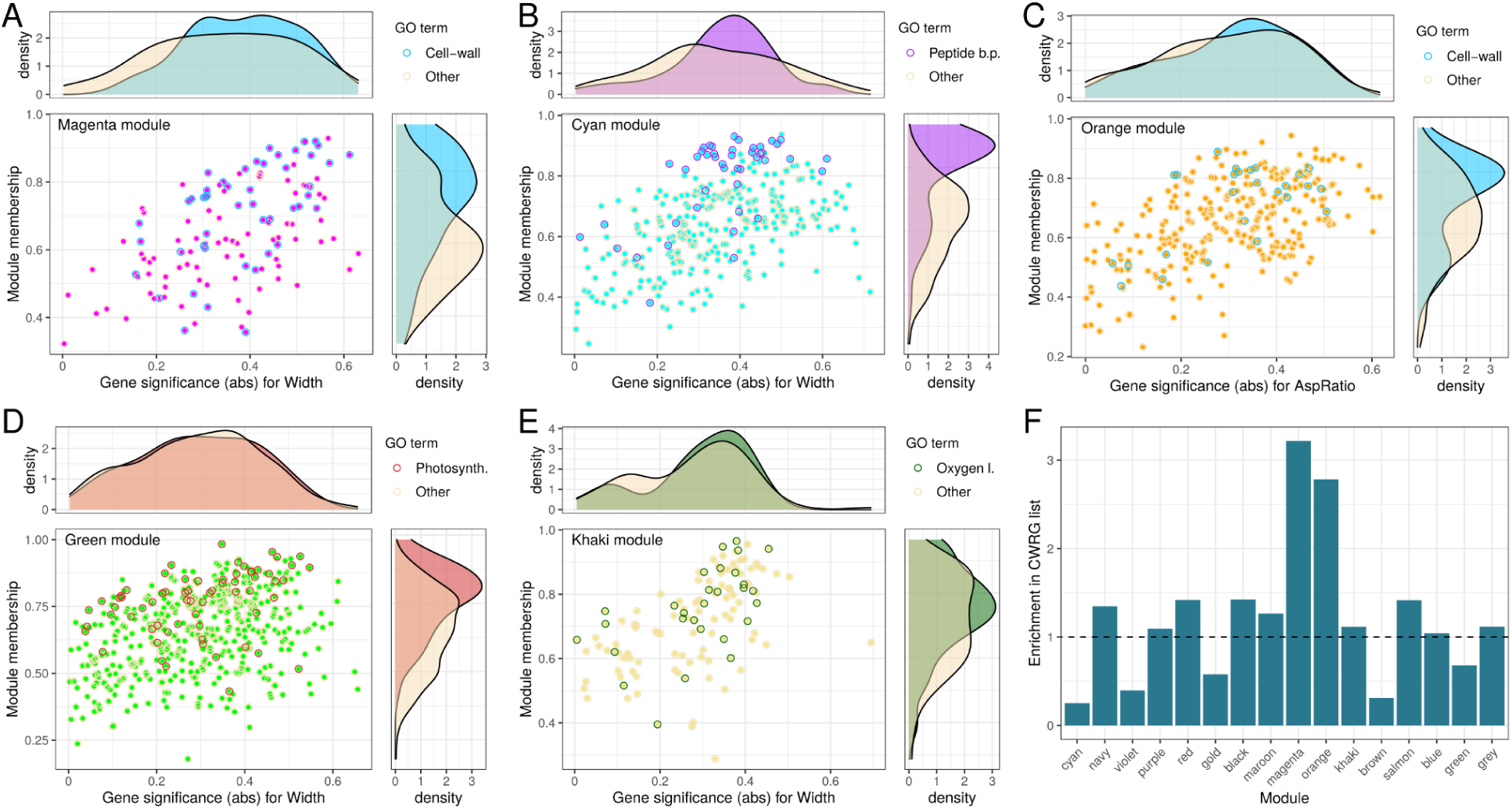
Relevance of genes associated with GO terms within each module. (A-E) Scatter plots showing module membership vs. gene significance (in absolute values and for the top-correlated parameter of the corresponding module) for all genes found within the magenta (A), cyan (B), orange (C), green (D), and khaki (E) modules. Highlighted points in each scatter plot correspond to genes associated with specific GO terms, which were the most enriched in each module: cell wall organization or biogenesis (A & C), peptide biosynthetic process (B), photosynthesis (D), and response to oxygen levels (E). Above each scatter plot, density plots for gene significance absolute values are shown for genes associated or not with the corresponding GO term. To the right of each scatter plot, density plots for module membership values are shown for genes associated or not with the corresponding GO term. (F) Enrichment in genes within our cell-wall related gene list across all modules. Enrichment values are calculated by dividing the observed number of CWRGs in each module by the expected number according to module size.

We further tested the conclusion that cell-wall related functions are important in gene modules that correlate with morphology. First, to assess the relevance of GO term enrichment results, we searched for members of our curated cell-wall related gene list across modules and we confirmed that the magenta and orange modules are enriched in cell-wall related genes (Figure 5F). Second, since response to stimulus and cell wall were GO terms that appeared enriched in highly variable genes and in modules most correlated with morphology, we verified that the latter enrichment was not merely a consequence of the former. We found that even though the khaki and magenta are the modules with highest CV^2^ of gene expression, the orange module does not have particularly high CV^2^ values and, in fact, both the cyan and green modules have low CV^2^ values (Supplementary Figure 7A). Furthermore, within the modules themselves, CV^2^ values do not correlate with either module membership or gene significance (Supplementary Figures 7B-7F). Altogether, this suggests that a strong correlation with morphology for cell-wall related gene modules is not an artifact of high variability in cell-wall related gene expression.

Additionally, cell-wall related genes with a high module membership within the magenta and orange modules are particularly well correlated between them, as if they were coregulated (Figure 6 & Supplementary Figure 8). We, hence, proceeded to reconstruct the GRN of cell-wall related genes within the magenta and orange modules, shown in Figure 6C & Supplementary Figure 8B, respectively, from their gene expression levels using GENIE3 (Huynh-Thu et al., 2010). No cell-wall related transcription factors are found within the orange module. However, within the magenta module we found two cell-wall related transcription factors with high module membership: *KNAT7* appears as a central node, and *NAC007* appears as a connector between two sub-networks. Both transcription factors are known to be involved in secondary cell-wall deposition (Nakano et al., 2015; Wang et al., 2020). *KNAT7* downstream targets identified by Li et al. (2012) involved in cellulose synthesis in secondary cell wall (*CESA4*, *CESA7* and *CESA8*) and xylan biosynthesis (*IRX8*/*GAUT12* and *IRX10*) are found in the magenta module and most are strongly connected in our GRN reconstruction (except for *CESA8*). *NAC007* (also known as *VND4*) has been described as playing a role in regulating secondary cell wall formation (Nakano et al., 2015), although its downstream targets have not been directly characterized, to our knowledge, by knockout mutants. There is, however, DAP-Seq data on *NAC007* (O’Malley et al., 2016). Using the ConnecTF tool (Brooks et al., 2021), we observed that out of the 3338 predicted targets of *NAC007*, 60 are within the magenta module, when only 30 would be expected if chosen at random across modules (log2 fold change of 0.98 and p-value of 1.59e-5), confirming that predicted targets of *NAC007* are enriched in the magenta module.

**Figure 6.**
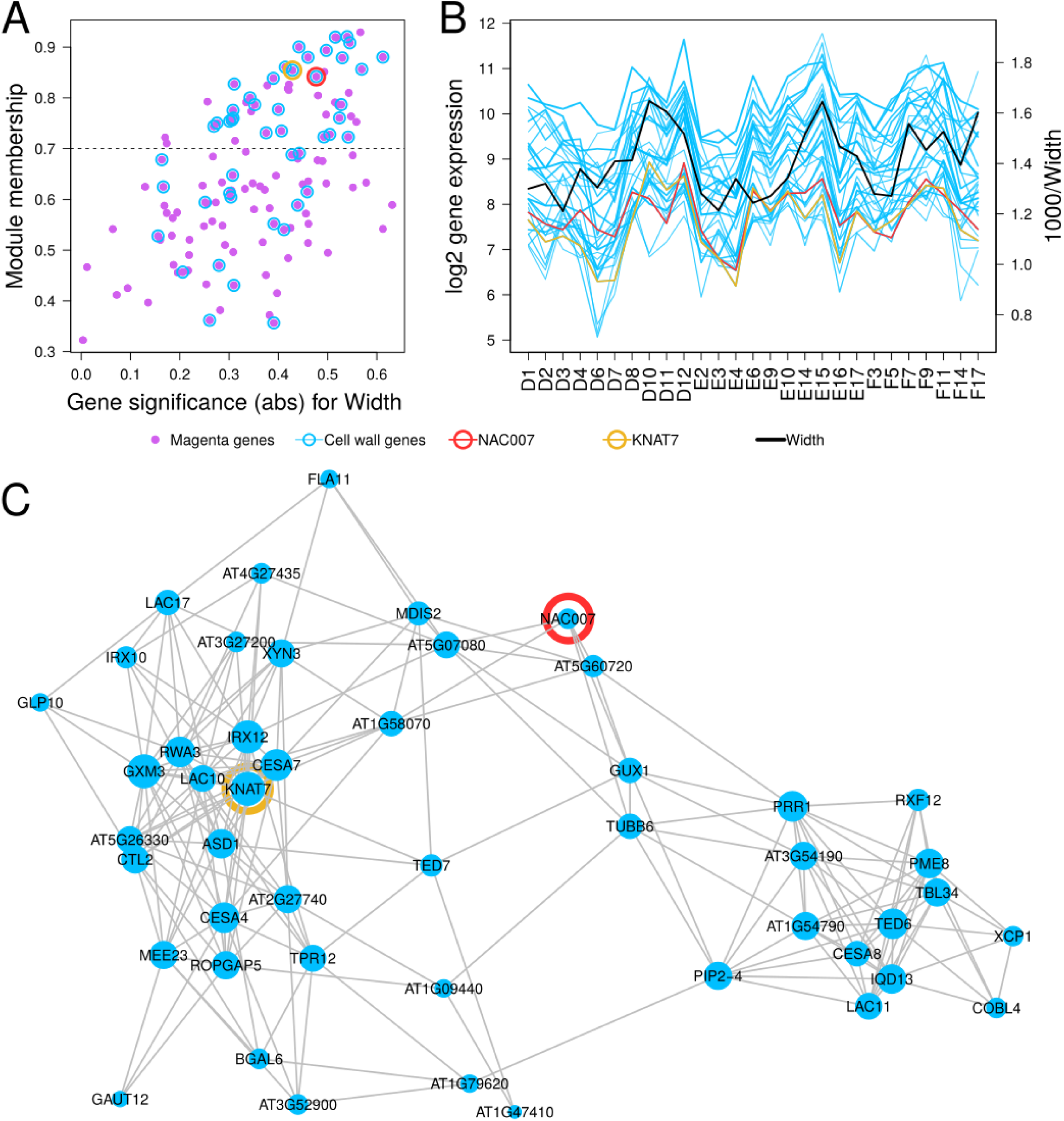
Properties of the magenta module and underlying gene regulatory network. (A) Module membership against gene significance in absolute value with respect to width for all genes from the magenta module. Higher module membership value indicates a gene that is more central in the module, while higher gene significance indicates that the expression of the gene has a higher correlation with width. Cell-wall genes are highlighted with blue rings, and genes encoding transcription factors *NAC007* and *KNAT7* with red and yellow rings, respectively. (B) Gene expression values across 27 samples for cell-wall related genes within the magenta module with module membership above 0.7 (black dotted line in A). Genes encoding transcription factors *NAC007* and *KNAT7* are shown in red and yellow, respectively. The black line corresponds to 1000/Width values for each sample (right axis), highlighting the negative correlation between gene expression of genes with high module membership in the magenta module and Width. The right axis has been chosen for Width measurements to overlap with gene expression values in order to visually show the negative correlation among them. (C) Gene regulatory network of all cell-wall related genes in the magenta module (see Methods/Gene regulatory network reconstruction). *NAC007* and *KNAT7* are highlighted in red and yellow, respectively. Node size is proportional to the degree (number of connections) of each node.

## Discussion

Sepals are important organs of angiosperm flowers. As with many other plant structures, and in particular floral organs, their morphology has been studied from a broad variety of approaches. We have here explored an approach that combines acquisition of sepal 3D images, automated extraction of relevant morphological parameters, exploration of natural variability in morphology and in gene expression across wild-type sepals, and reconstruction of gene regulatory networks from correlations between gene expression patterns and morphology. In doing so, we have found cell-wall related genes as having highly variable expression in wild-type sepals. Furthermore, gene regulatory networks that correlate with morphology are associated with cell wall organization and biogenesis.

We consider that our approach is complementary to other approaches, such as mutant or condition-based, used to detect genes affecting morphology. In a classical mutant approach, wild-type variability is explored primarily as a means to control for different mutant conditions across biological replicates. Instead, we here make use of such variability to extract biologically relevant information. In particular, we use variability in gene expression and morphology in wild-type plants of the same accession to unravel gene regulatory networks correlating with organ morphology. To recover modules of co-expressed genes, we used the WGCNA package (Langfelder & Horvath, 2008) which requires relatively few parameters to adjust. Other tools, such as sPLS (Lê Cao et al., 2008), also allow for the association of two-block data, like gene expression and morphology. However, sPLS recovers individual genes instead of modules. Given the risk of false positives when focusing on small subsets of genes, we considered that associating morphology to modules of co-expressed genes instead of individual genes was a more conservative approach.

Based on our morphology analysis, we predicted that area and aspect ratio were the most informative parameters to describe sepal size and shape. However, our WGCNA results show width as the most highly correlated parameter. Nevertheless, when viewing all statistically significant correlations, we observe that modules that correlate with width also correlate with area (magenta and cyan modules) and that the third most significantly correlated module (orange) correlates with aspect ratio, which fits in well with the aforementioned prediction. Looking at enriched GO terms for each module (Supplementary Table 5) we find that two modules are strongly enriched in cell-wall related genes (magenta and orange), and another one (cyan), in chloroplast-related functions and peptide biosynthesis. The cell wall is known to strengthen the plant body and to play a key role in plant growth. During evolution, plant cells have acquired the capacity to synthesize walls made of polysaccharides, to assemble them into a strong fibrous network and to regulate cell-wall expansion during growth (reviewed in Cosgrove, 2005). The polysaccharides that contribute to the biomass and to the cell wall are made up of sugars produced in chloroplasts, as end products of photosynthesis. Finding cell-wall and chloroplast related genes associated with sepal morphology is therefore consistent with known biological processes.

The contribution of different sets of co-regulated genes to sepal morphology can be thought of in isolation but connections between modules can also be established, albeit with a certain degree of speculation. For example, the magenta module correlates negatively with width, whereas the cyan module correlates positively with width. One could speculate that genes in the magenta module could have repressive effects on growth, i.e., secondary cell wall might be involved in the growth repression; and that the genes in the cyan module could have a contrary effect, i.e., increased peptide biosynthesis associated with increased growth. Additionally, the fact that the orange module, also enriched in cell-wall related genes, correlates best with aspect ratio and not with width like the magenta module, could indicate that different mechanisms are at work to control size and shape. In fact, the alignment of cortical microtubules is important for the anisotropy of sepal growth and in determining the aspect ratio of mature sepals (Hervieux et al., 2016). Interestingly, supramolecular fiber and cytoskeleton organization genes appear to be overrepresented in the orange module.

Whereas other studies have aimed at reconstructing complete and consistent gene regulatory networks based on reviews of the literature (e.g. Espinosa-Soto et al., 2004; La Rota et al., 2011), we have shown that variability of gene expression in wild type allows the recovery of gene sets that likely function together. An illustration that this is likely the case, is that we find two transcription factors which are known to dimerize, BZIP34 and BZIP61, in the orange module. If we focus on the magenta module, which contains genes related to the cell wall, we find that the top two transcription factors of this module (ordered by module membership), NAC007 and KNAT7, are known to be involved in secondary cell-wall deposition and to belong to the same GRN (Wang et al., 2020). As expected, the identified direct targets of NAC007 (O’Malley et al., 2016) are enriched in the magenta module. Although DAP-Seq data is not currently available for KNAT7, some known targets of KNAT7, such as *IRX8/GAUT12* and *IRX10* (Li et al., 2012; He et al., 2018), belong to the magenta module, as shown in Figure 6C. In addition, three *CESA* genes that have been shown to be upregulated in the *knat7* loss-of-function mutant (Li et al., 2012) also belong to the magenta module. Interestingly, the *knat7* mutant has reduced seed size (Renard et al., 2020) and a repressor version of KNAT7 (supposed to mimic the mutant) induces a dwarf phenotype (Qin et al., 2020), supporting a link between KNAT7 and growth control. A speculative mechanism could be that KNAT7 positively regulates secondary cell-wall deposition, as shown by Wang et al. (2020), which is known to rigidify the tissues (Zhong & Ye, 2015) and could potentially lead to growth arrest. This is consistent with the magenta module being negatively correlated with width and area.

Despite having grown our three plants in experimentally controlled conditions and all of them having grown healthy, we observe that genes involved in response to stimulus are highly variable among our sepal samples. Although we cannot completely exclude the possibility that expression of response to stimulus genes was induced by sepal dissection and imaging, it is unlikely since we managed to keep the duration of sepal manipulation below five minutes. Our interpretation, in line with other studies (Araújo et al., 2017, Cortijo et al., 2019), is that this variability is not in itself due to differences in gene expression as a response to strong external stimuli experienced by different sepals, but rather, the evidence of underlying basal gene expression variability allowing the plant to cope with small microenvironmental differences. This basal variability is thought to have been selected to allow for the possibility of adaptation to environmental change (Queitsch et al., 2002). We extend this interpretation to cell-wall related genes, which also show significant variability of expression in sepals. Considering that reproducibility in morphology should be achieved in part thanks to underlying regulatory and compensatory mechanisms, our results indicate that cell-wall related genes could be fundamental for these mechanisms. The possible role of cell-wall related genes in determining sepal morphology could be applicable to other organs since cell-wall related genes exhibit high variability across plants also in Arabidopsis seedlings (Cortijo et al., 2019). In general, our work sheds light on the links between expression variability, gene regulatory networks, and developmental robustness and thereby, opens the way to informed functional analysis.

## Materials and Methods

### Plant Material

Sibling Col-0 plants were grown on soil at 20°C in short day conditions (8 h light/16 h dark) for 20 days before being transferred to long day conditions at 22°C (16 h light/8 h darkness). Buds were dissected from independent secondary inflorescences after at least 10 siliques were formed and were observed under the binocular to identify those at the beginning of stage 11 (stigmatic papillae formed, petals being longer than the short stamen but shorter than the long stamen) as described by Smyth et al. (1990). After dissection, each abaxial sepal was transferred to 0.8% agarose to avoid dehydration, imaged with a confocal microscope and immediately transferred to liquid nitrogen. The manipulation and the confocal imaging lasted less than 5 min to minimize the impact of the manipulation on the transcriptome.

### Confocal Imaging

Sepals were examined in a Leica SP5 confocal microscope equipped with a 10X objective. Samples were imaged using laser 488 nm set up at 20% of the maximum power and emission from 498 to 735 nm was recovered. The acquisition time was no more than 100 seconds.

### Segmentation and extraction of geometrical parameters

Images are pretreated in the xy plane using a Gaussian filter of sigma of the order of the z direction voxel size. Then a normalizing procedure is performed on the whole filtered sample. The linear normalization is constructed in such a way that it superimposes the background value and the Otsu threshold value of the output images. The sepal’s contour detection procedure is based on the EdgeDetect Morphology process of the MorphoGraphX software. First, a rough contour is assessed as the intersection of the top-down and the bottom-up contours given by Edge Detect. Then a dilation in the xy direction is performed in order to recover the weakly marked margin cells of the sepals. The principal directions of the sepal contour are computed, and sepals are placed in the frame where their center of mass is in the origin (their first principal axis is along the Oy axis and the second principal axis is along the Ox axis). The length measurements are done on the longitudinal and transversal principal sections using a python script. The sepal’s abaxial and adaxial surfaces are separated by watershed segmentation of the maximal curvature map constructed on the sepal’s whole surface. Then the area of the abaxial surface is computed as areas of cells on a tissue surface mesh in MorphoGraphX. The morphological parameters measured are length, width, area, longitudinal curvature and transversal curvature. Aspect ratio is length divided by width and Gaussian curvature is the product of longitudinal and transversal curvatures.

In order to validate geometrical measurements, we compared a sample of 10 images from 10 different sepals with a sample of 10 images of the same sepal that was manipulated between consecutive shootings. For each of our parameters, same-sepals variability was verified to be less than 5% of the variability measured across different sepals. Other parameters that had originally been measured, such as curvature at the sepal periphery, were excluded because they did not pass this validation criterion.

### RNA extraction and sequencing

Each sepal was ground individually using a plastic conical pestle fitting the bottom of the Eppendorf tube, which was dipped in liquid nitrogen. RNA extraction was subsequently performed using the Arcturus PicoPure RNA Isolation Kit from Thermo Fisher Scientific following the manufacturer’s instructions. The libraries were sequenced by the HELIXIO company on Illumina NextSeq 500 using single read sequencing of 76bp in length.

### Quantification of gene expression

Raw reads were pseudo-aligned to cDNA and non-coding RNA sequences from the TAIR10 *Arabidopsis thaliana* reference genome (Ensembl release 47) using kallisto (Bray et al., 2016). Transcript abundances obtained were converted to gene abundances using tximport (Soneson et al., 2015). Gene counts per million were obtained using the edgeR package (Robinson et al., 2010) in R (R Core Team, 2021). Only genes with at least 5 counts per million in at least 14 out of 27 sepal samples were retained and counts per million were normalized using the a trimmed mean of M-values (TMM) method in edgeR, resulting in 14,085 gene expression values for 14,085 genes. We validated our bulk RNA-Seq results by comparing them to RNA-Seq samples obtained from Arabidopsis seedlings from Cortijo et al. (2019) (Supplementary Figure 9).

### WGCNA

Parameters used for WGCNA were: soft-thresholding power for adjacency matrix reconstruction, 11; minimum module size, 50; module merging threshold, 0.35. Each gene within a module has a module membership value, calculated as the correlation between that gene’s expression and the expression of the module eigengene across all samples. It is representative of the gene’s intramodular connectivity. Each gene also has a gene significance for any trait, calculated as the correlation between the gene’s expression values and the morphological parameter values across all samples.

### GO enrichment analysis

Gene ontology enrichment analysis was performed with PANTHER17.0 (Mi et al., 2021) at http://pantherdb.org (last accessed on 2022-03-30) with the following parameters: Analysis Type: PANTHER Overrepresentation Test (Released 2022-02-02); Annotation Version and Release Date: GO Ontology database doi:10.5281/zenodo.5725227 Released 2020-11-01; Test Type: FISHER; Correction: FDR. Background used was our 14,085 gene dataset (Supplementary Table 5), and Arabidopsis whole genome (Supplementary Table 6). Top hierarchy GO terms were obtained by filtering the PANTHER output with the following criteria: only positive enrichment terms were kept, only top hierarchy terms in each block (as output by PANTHER) were kept according to total number of genes associated with each GO term, terms were ordered according to their false discovery rate (FDR) starting from the smallest rate, only the top five resulting terms were kept for each highly-variable group (HVG) or lowly-variable group (LVG) group unless a term in the top five in one group also appeared in another group, in which case it was kept in both. Log2 fold change values (Supplementary Table 7) and FDR (Supplementary Table 8) were calculated for each of these terms in each HVG or LVG group.

### Cell-wall related gene list

To produce this list, first, we referred to a review describing all the enzymes acting on the cell wall (Frankova and Fry, 2013) and identified the corresponding genes in the TAIR Arabidopsis database (www.tair.com). Second, we used the CAZy database (www.cazy.org) describing families of structurally-related catalytic and carbohydrate-binding modules of enzymes that degrade, modify or create glycosidic bonds to find corresponding genes in the Arabidopsis genome. Finally, we manually enriched the list with proteins known to directly interact with those enzymes in the cell wall, such as pectin methylesterase inhibitors or interactors of cellulose synthase, also adding the extensins, which are non-enzymatic proteins highly abundant in the cell wall and important for its biosynthesis.

### Gene regulatory networks

Gene regulatory network reconstruction was performed with GENIE3 (Huynh-Thu et al., 2010) with the Random Forest method and default parameters. The weighted adjacency matrix obtained was modified by setting all adjacent values above 0.045 to 1 and below 0.045 to 0. The 0.045 threshold was chosen to improve visibility and to include all cell-wall related genes in the modules. Graph visualization was performed by using the graph_from_adjacenty_matrix function from the igraph (version 1.2.11) package and the ggnet2 function in R (R Core Team, 2021). Genes under the GO term “cell wall organization or biogenesis” (GO:0071554) were extracted from Agrigo2.0 (Tian et al., 2017) at http://systemsbiology.cpolar.cn/agriGOv2/ (last accessed on 2023-02-08) and are listed in Supplementary Table 9.

### Variability of gene expression

Squared coefficient of variation (CV^2^) of gene expression values were calculated by dividing the variance over the square of the mean of gene expression measurements across 27 sepal samples.

### Data, script and code availability

Supplementary Tables 1-9, R scripts (R Core Team, 2021) used for analysis and to generate main and supplementary figures, as well as all data necessary for analyses are available at https://doi.org/10.5281/zenodo.8146786. Image analysis code and pipeline for morphological parameter extraction is available at http://forge.cbp.ens-lyon.fr/redmine/projects/florivar. Raw sepal image data are available at https://doi.org/10.5281/zenodo.6559804. Raw RNA-Seq data for 30 sepal samples are available at the European Nucleotide Archive (ENA) under project PRJEB52917.

## Acknowledgements

We wish to acknowledge S. Cortijo for valuable suggestions in the review process, A. Roeder for constructive discussion and for suggestions on a first version of the manuscript, A. Ahmed for work on a pilot project, PLATIM for provision of microscope facilities, Helixio for their excellent service and assistance in the RNA sequencing, J. Pelloux for sharing mutants and discussion on cell wall, Y. Long, P. Das, A. Fruleux and S. Bovio for comments and suggestions, and A. Lacroix, J. Berger, P. Bolland, H. Leyral and I. Desbouchages for assistance with plant growth and logistics.

## Funding

This work was funded by the French National Research Agency (ANR) through a European ERA-NET Coordinating Action in Plant Sciences (ERA-CAPS) grant (Grant No. ANR-17-CAPS-0002-01 V-Morph) and through a direct grant (Grant No. ANR-17-CE20-0023-02 WALLMIME), by Fond de Recherche ENS Lyon (Projet émergent FLORIVAR), and by BAP INRAE (Projet FLORIVAR).

## Author contributions

Conceptualization: DAH, AB, FMo

Methodology: DAH, AK, FMa

Software: DAH, AK, MBV

Validation: DAH, VB, CM

Formal Analysis: DAH, AK

Investigation: DAH, AK, VB, CS, ADV, FMa, MMM, FMo

Data Curation: DAH, AK, VB, CS, MBV, FMo

Writing - Original Draft Preparation: DAH, AB, FMo

Writing - Review & Editing: DAH, AK, MMM, AB, FMo

Visualization: DAH, AK, MBV, FMo

Supervision: DAH, AB, FMo

Project Administration: AB, FMo

Funding Acquisition: AB, FMo

## Conflict of interest disclosure

The authors declare they have no conflict of interest relating to the content of this article.

---

**Supplementary Figure 1:**
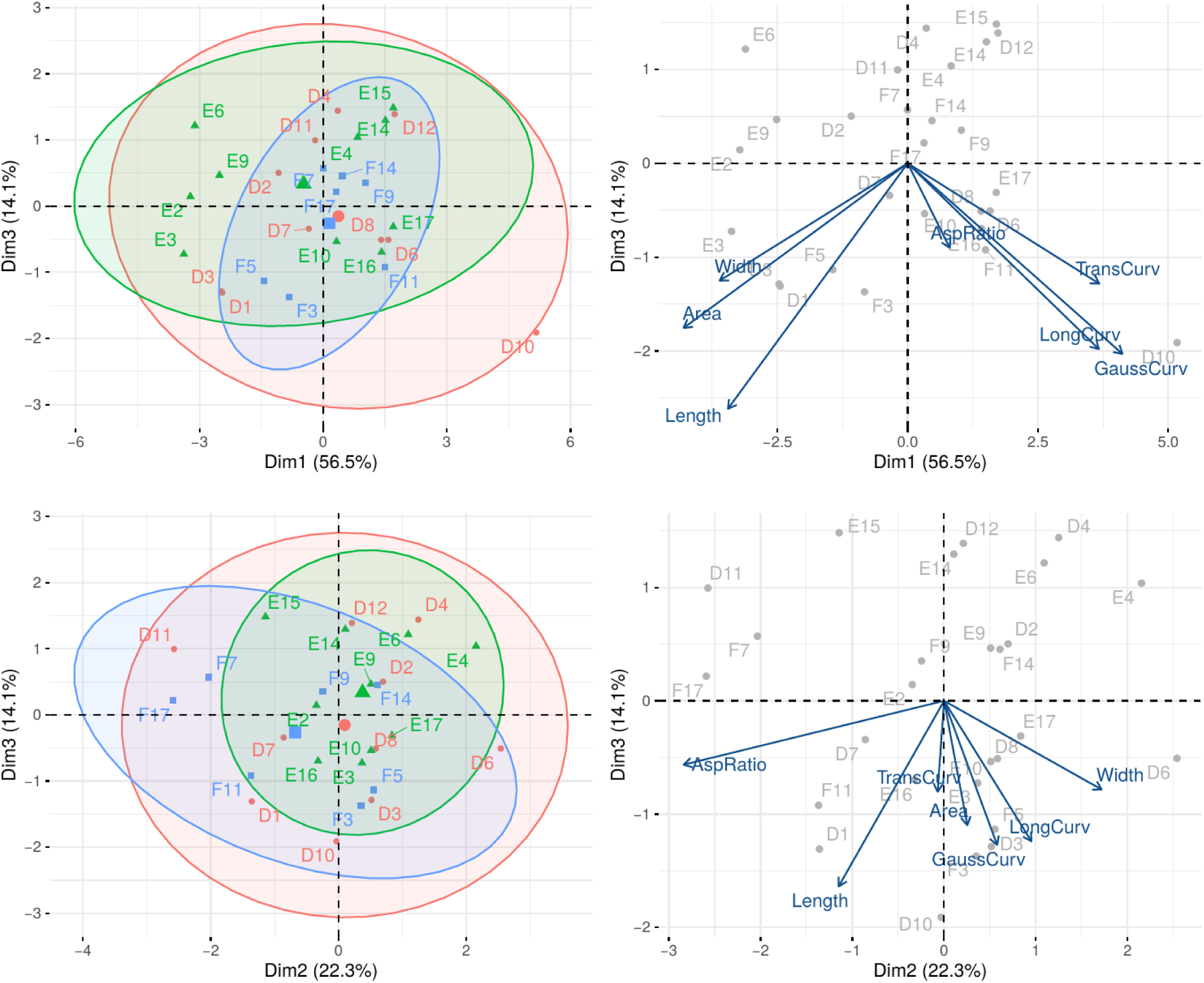
Principal component analysis (PCA) for all samples according to their morphological parameters (plant D in red, E in green, F in blue), showing the percentage of variance explained by each principal component. Top: PC1 vs. PC3. Bottom: PC2 vs. PC3. Left: shaded areas show 95% confidence ellipses for each set of 2D normally distributed samples (plant D in red, E in green, F in blue). Right: loadings for the PCA depicting the contribution of each morphological parameter to each principal component in the 2D space.

**Supplementary Figure 2:**
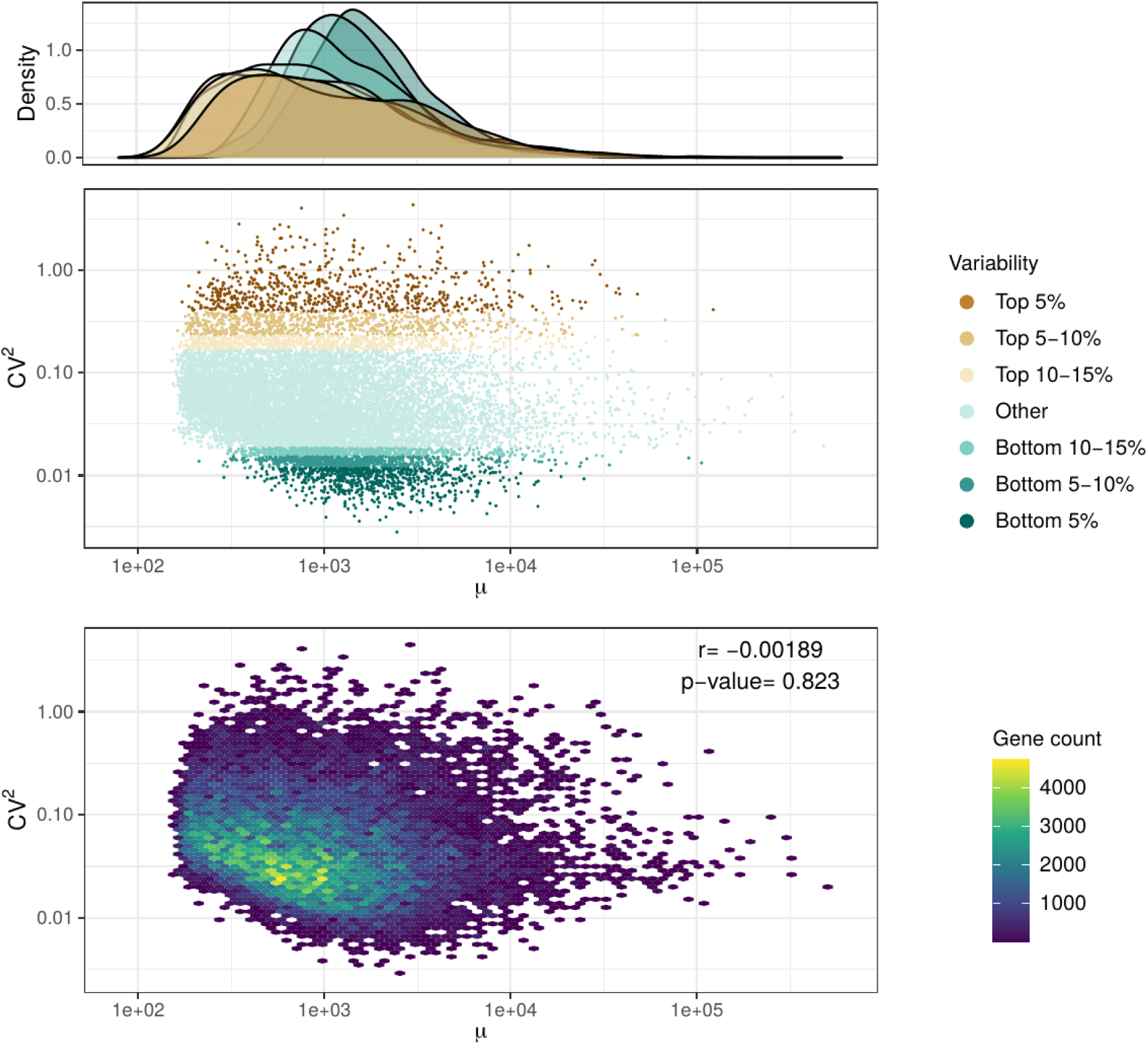
Squared coefficient of variation (CV^2^) against average (μ) of gene expression (middle plot) across 27 samples for 14,085 genes identifying highly variable genes (top 5%, top 5-10%, and top 10-15%) and lowly variable genes (bottom 5%, bottom 5-10%, and bottom 10-15%). Density plots for each CV^2^ classification are shown on top. Point density in the 2 dimensional space (CV^2^ against μ) is shown in the bottom plot with the Pearson correlation coefficient (r) and the corresponding p-value shown highlighting no correlation between the two variables.

**Supplementary Figure 3:**
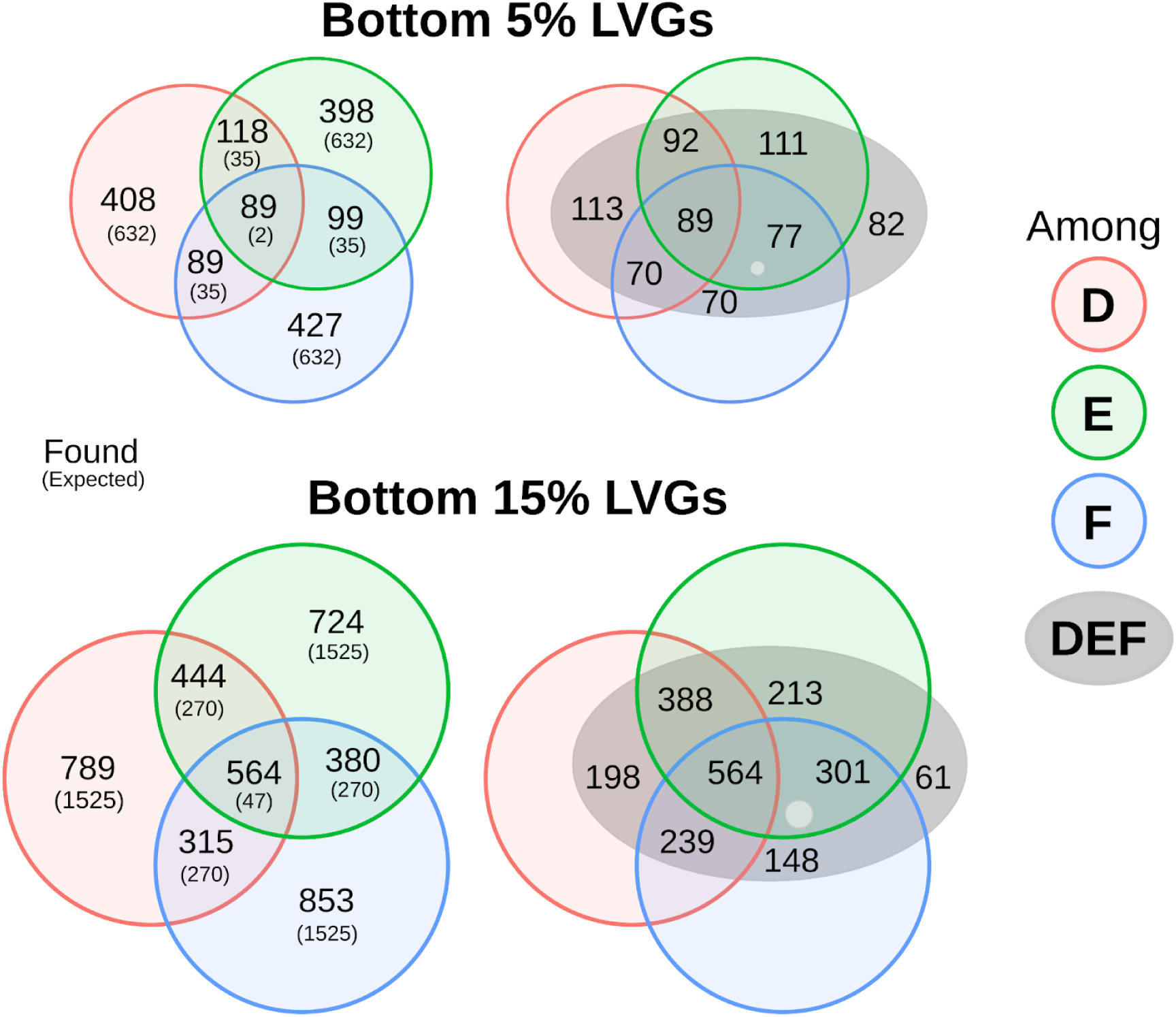
Venn diagrams for the bottom 5% lowly variable genes (LVGs) and for the bottom 15% LVGs according to their gene expression CV^2^ values. Lowly variable genes were extracted in two different ways: considering sepals from plants D & E & F independently (left), and considering variability among 27 sepals from all three plants pooled together (DEF; right). The left Venn diagrams show the overlap between genes found for each plant independently. In the top figures, the total number of genes in each circle (red, green and blue) is 704, corresponding to 5% of our complete 14,085 gene set. Similarly, the bottom figures contain 2,112 genes in each circle (15% of 14,085). Numbers shown in parentheses are the expected number of common genes found in each intersection if genes had been chosen randomly for each plant. The right Venn diagrams show how the LVG set obtained by considering all 27 sepal samples (DEF) compares to the lowly variable gene sets obtained using each plant independently. Genes within the grey ellipse also sum up to 704 (top) and 2,112 (bottom). All genes in the intersection between the three plants are also present within the grey ellipse (89 out of 89, top; 564 out of 564, bottom), whereas only a few genes (82, top; 61, bottom) are found in the DEF set and not for any plant independently. Circle and ellipse sizes and intersections are drawn to aid visualization but their sizes are not proportional to the number of genes found within them. “Holes” in the grey ellipse correspond to genes that belong to the corresponding area in the left plot, but are not found within our DEF LVG set.

**Supplementary Figure 4:**
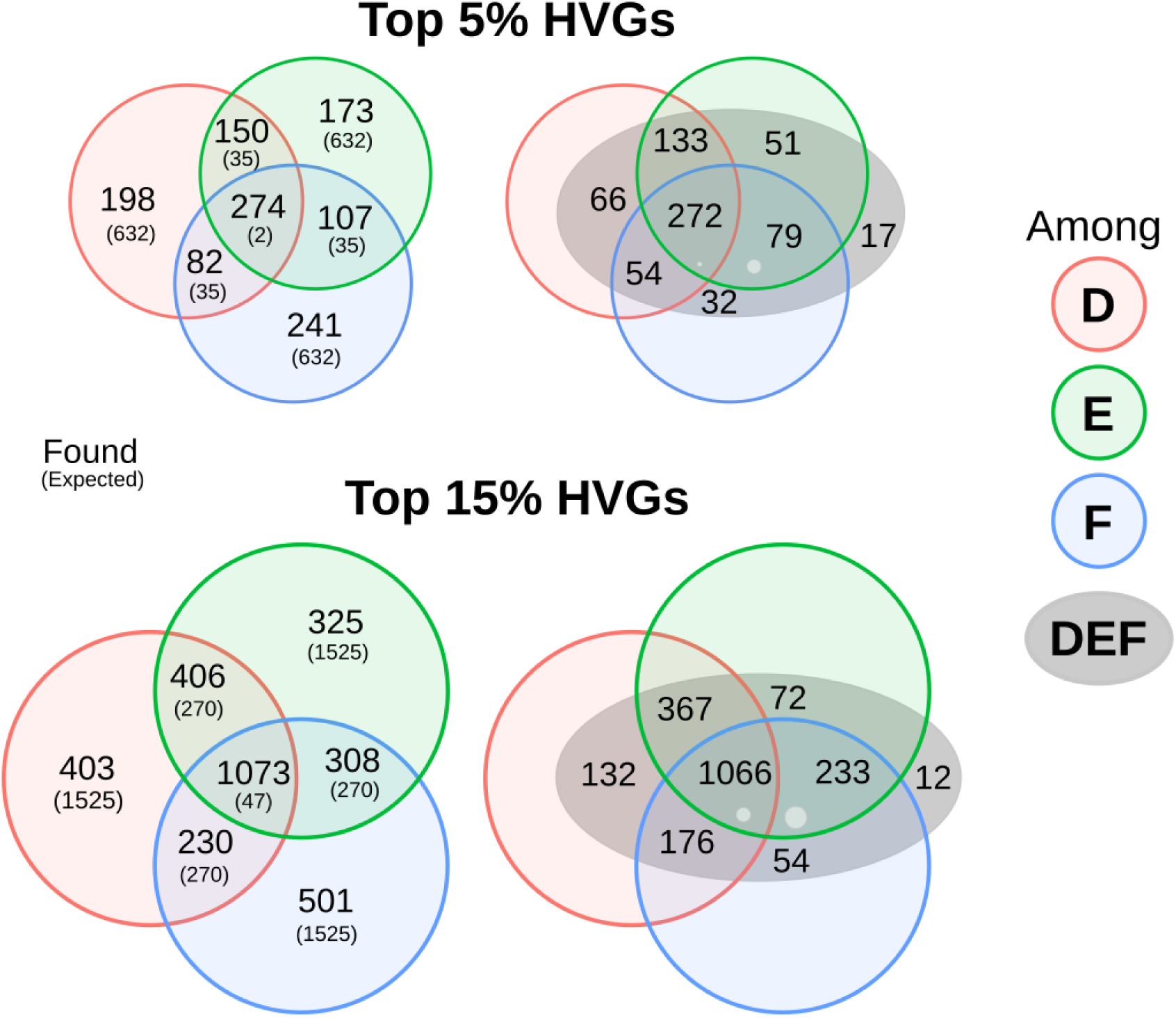
Venn diagrams for the top 5% highly variable genes (HVGs) and for the top 15% HVGs according to their gene expression CV^2^ values. Highly variable genes were extracted in two different ways: considering sepals from plants D & E & F independently (left), and considering variability among 27 sepals from all three plants pooled together (DEF; right). The left Venn diagrams show the overlap between genes found for each plant independently. In the top figures, the total number of genes in each circle (red, green and blue) is 704, corresponding to 5% of our complete 14,085 gene set. Similarly, the bottom figures contain 2,112 genes in each circle (15% of 14,085). Numbers shown in parentheses are the expected number of common genes found in each intersection if genes had been chosen randomly for each plant. The right Venn diagrams show how the HVG set obtained by considering all 27 sepal samples (DEF) compares to the highly variable gene sets obtained using each plant independently. Genes within the grey ellipse also sum up to 704 (top) and 2,112 (bottom). Almost all genes in the intersection between the three plants are also present within the grey ellipse (272 out of 274, top; 1066 out of 1073, bottom), whereas only a very small number of genes (17, top; 12, bottom) are found in the DEF set and not for any plant independently. Circle and ellipse sizes and intersections are drawn to aid visualization but their sizes are not proportional to the number of genes found within them. “Holes” in the grey ellipse correspond to genes that belong to the corresponding area in the left plot, but are not found within our DEF HVG set.

**Supplementary Figure 5:**
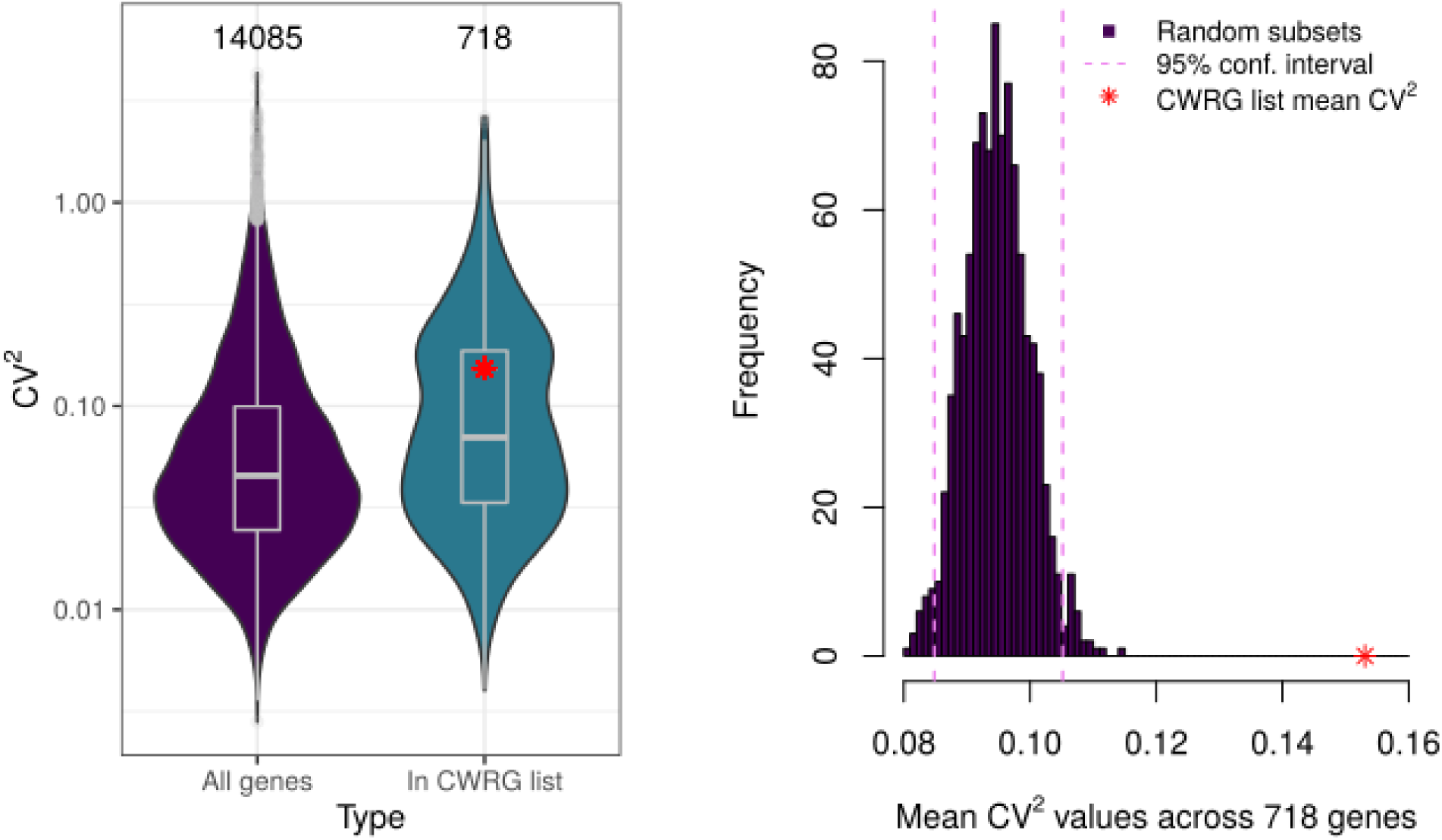
Left: violin plots of gene expression CV^2^ values for our entire 14,085-gene set (purple) compared to that of the 718 cell-wall related gene (CWRG) list found in the dataset (blue). Grey box plots show median, 1st and 3rd quartiles of the distributions. Right: histogram of gene expression CV^2^ mean values from 1000 random subsets of 718 genes from the 14,085-gene set. Pink vertical dashed lines delimit the 95% confidence interval of the distribution. Red asterisk shows the gene expression CV^2^ mean value of the CWRG list, clearly above the 95% confidence interval of the random subset gene expression CV^2^ distribution.

**Supplementary Figure 6:**
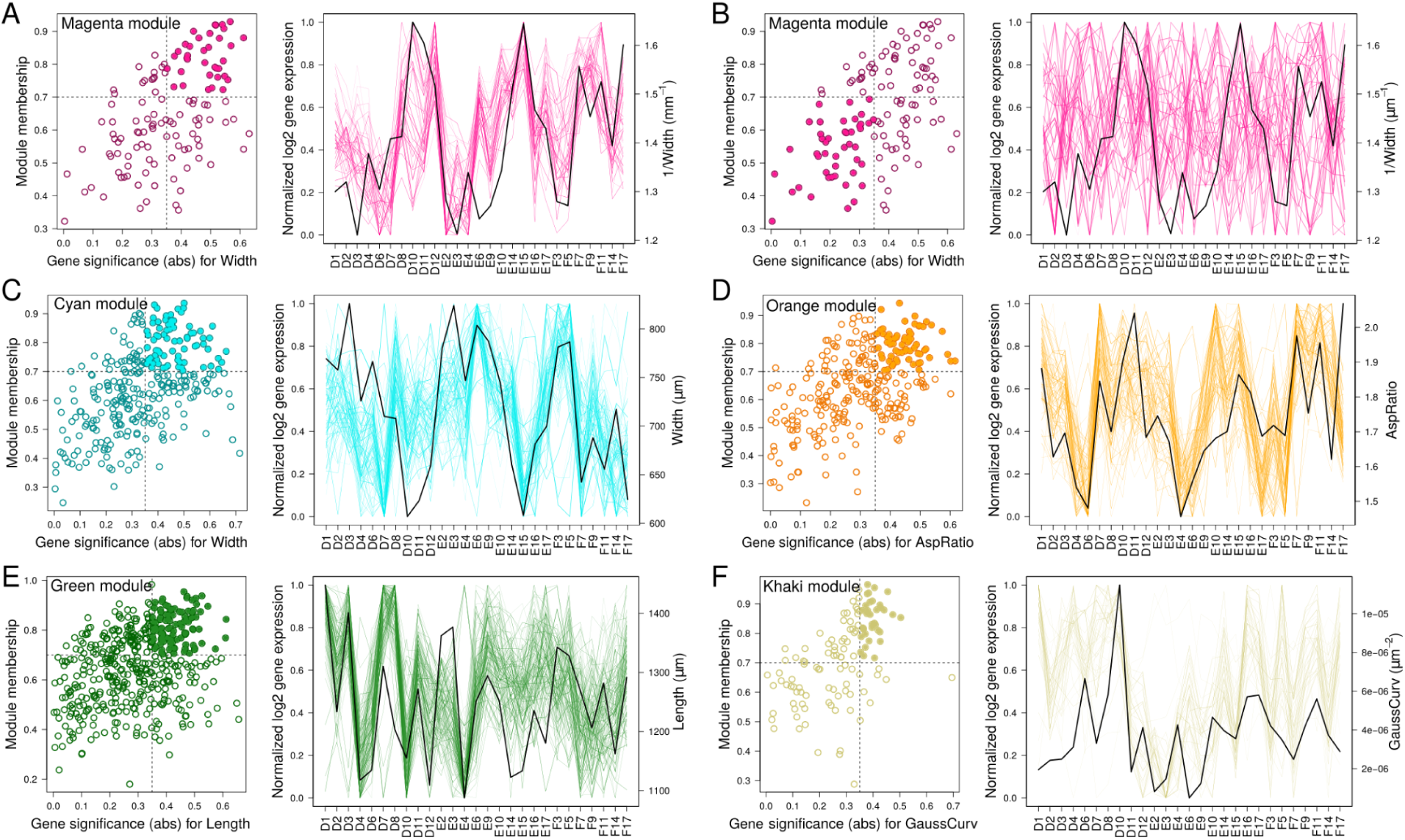
Module membership against gene significance for the top correlated morphological parameter of the corresponding module (left) and normalized (with 0 being the lowest expression and 1 being the highest expression across samples) log2 gene expression (right) values for 27 samples, for magenta (A & B), cyan (C), orange (D), green (E) and khaki (F) modules. Gene expression values are compared with the corresponding morphological parameter with a scale chosen so as to depict high correlation with gene expression for the highlighted genes. Only the genes with module membership above (below for B) 0.7 and gene significance above (below for B) 0.35 for the corresponding parameter, highlighted with filled circles (left plot), are plotted in the gene expression plot (right plot).

**Supplementary Figure 7:**
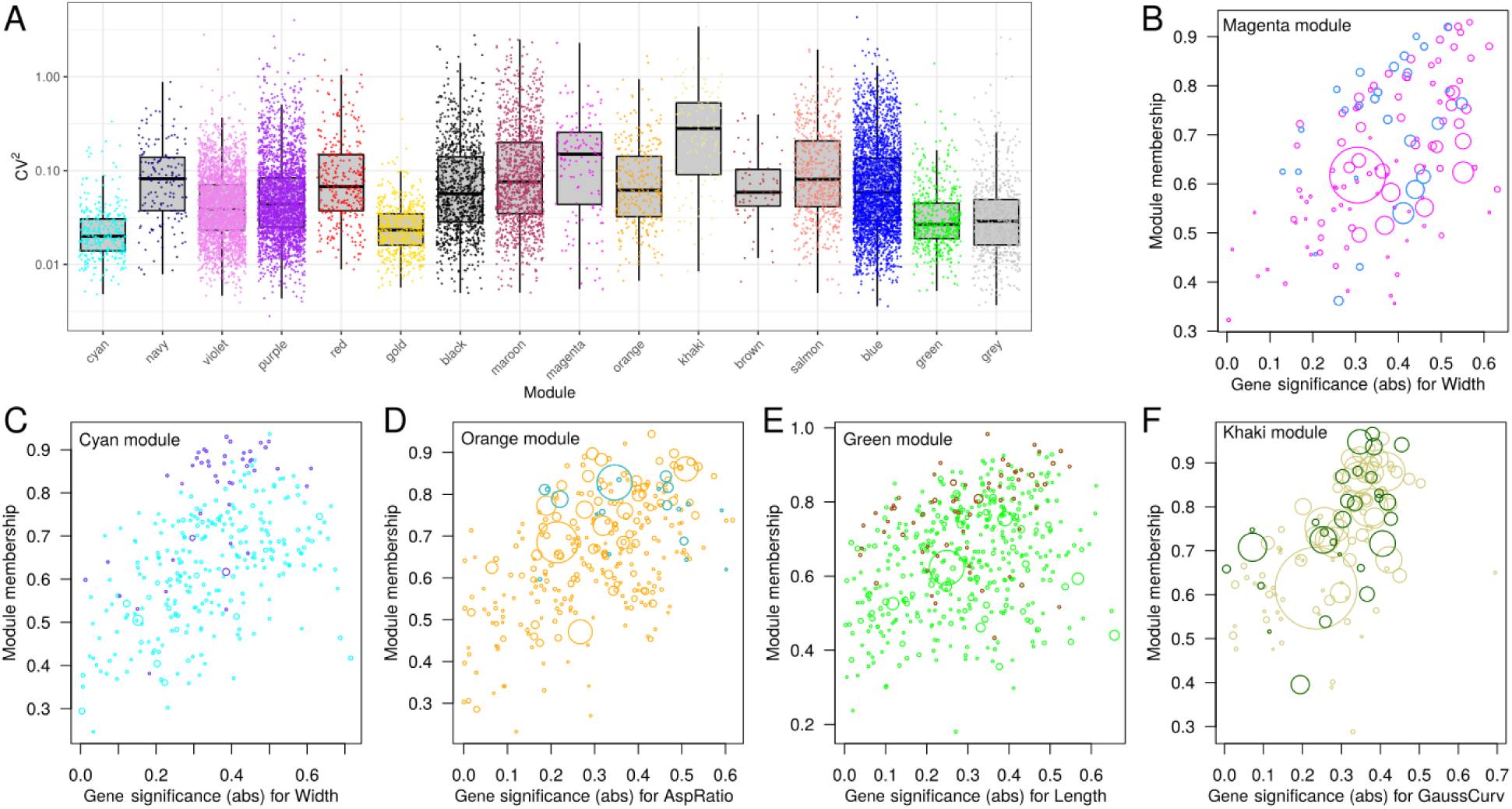
(A) Boxplots for CV^2^ of gene expression values for all genes within each module. Each color point represents one gene and the black bar in each boxplot represents the median value across all the genes in each module. (B-F) Scatter plots showing module membership vs. gene significance (in absolute values and for the top-correlated parameter of the corresponding module) for all genes found within the magenta (B), cyan (C), orange (D), green (E), and khaki (F) modules. Circle size is proportional to each gene’s CV^2^ value. Genes with a different color to the corresponding module name are those associated with the specific GO terms which were the most enriched in each module: cell wall organization or biogenesis (B & D), peptide biosynthetic process (C), photosynthesis (E), and response to oxygen levels (F).

**Supplementary Figure 8:**
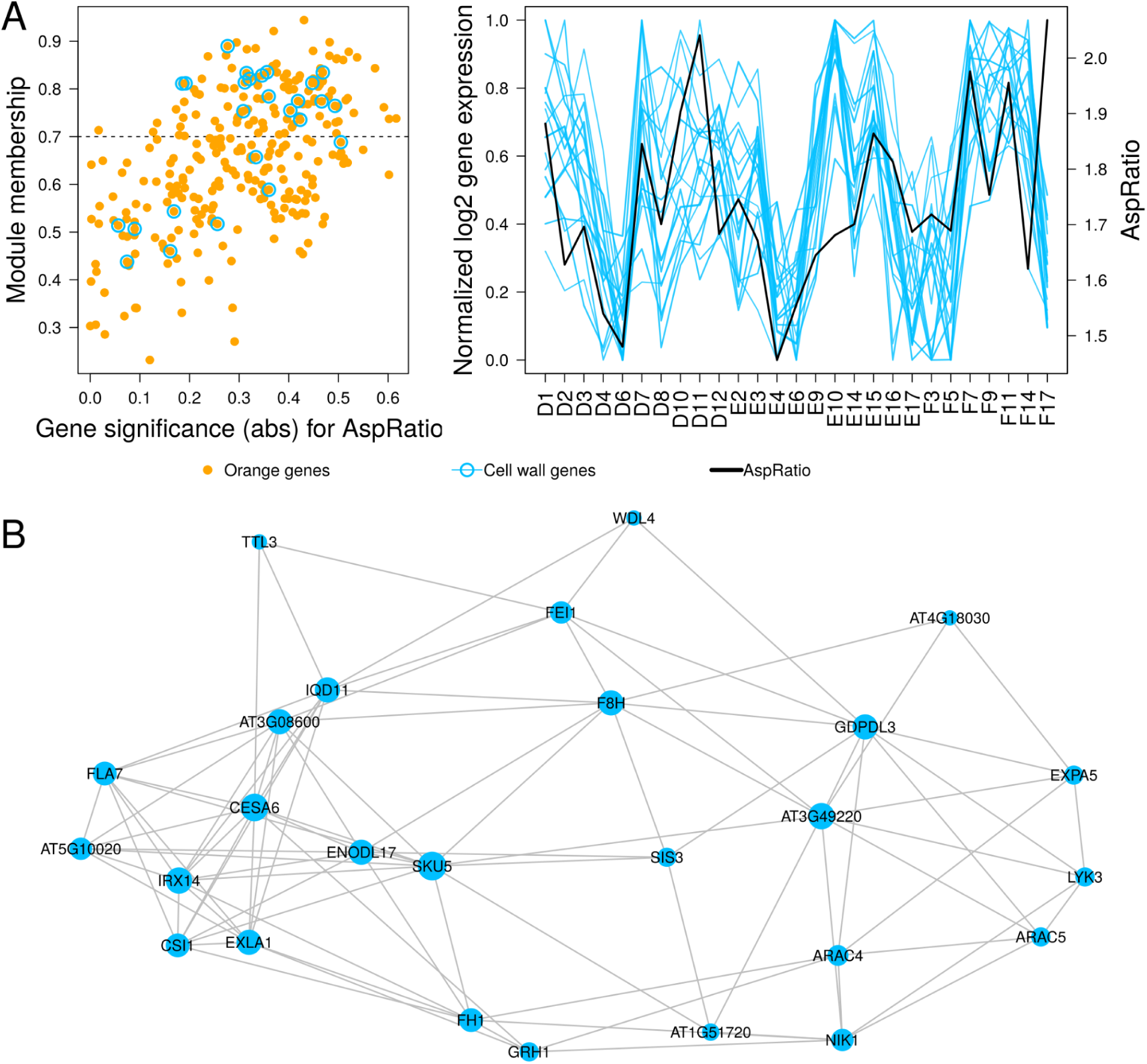
(A) Module membership against gene significance in absolute value with respect to AspRatio for all genes from the orange module are shown in the left plot. Cell-wall related genes are highlighted with blue rings and their normalized gene expression values across 27 samples (for those genes with module membership above 0.7) are shown in the right plot. The black line corresponds to AspRatio values for each sample (right axis). (B) Gene regulatory network of all cell-wall related genes in the orange module (see Materials and Methods). Node size is proportional to the degree (number of connections) of each node.

**Supplementary Figure 9:**
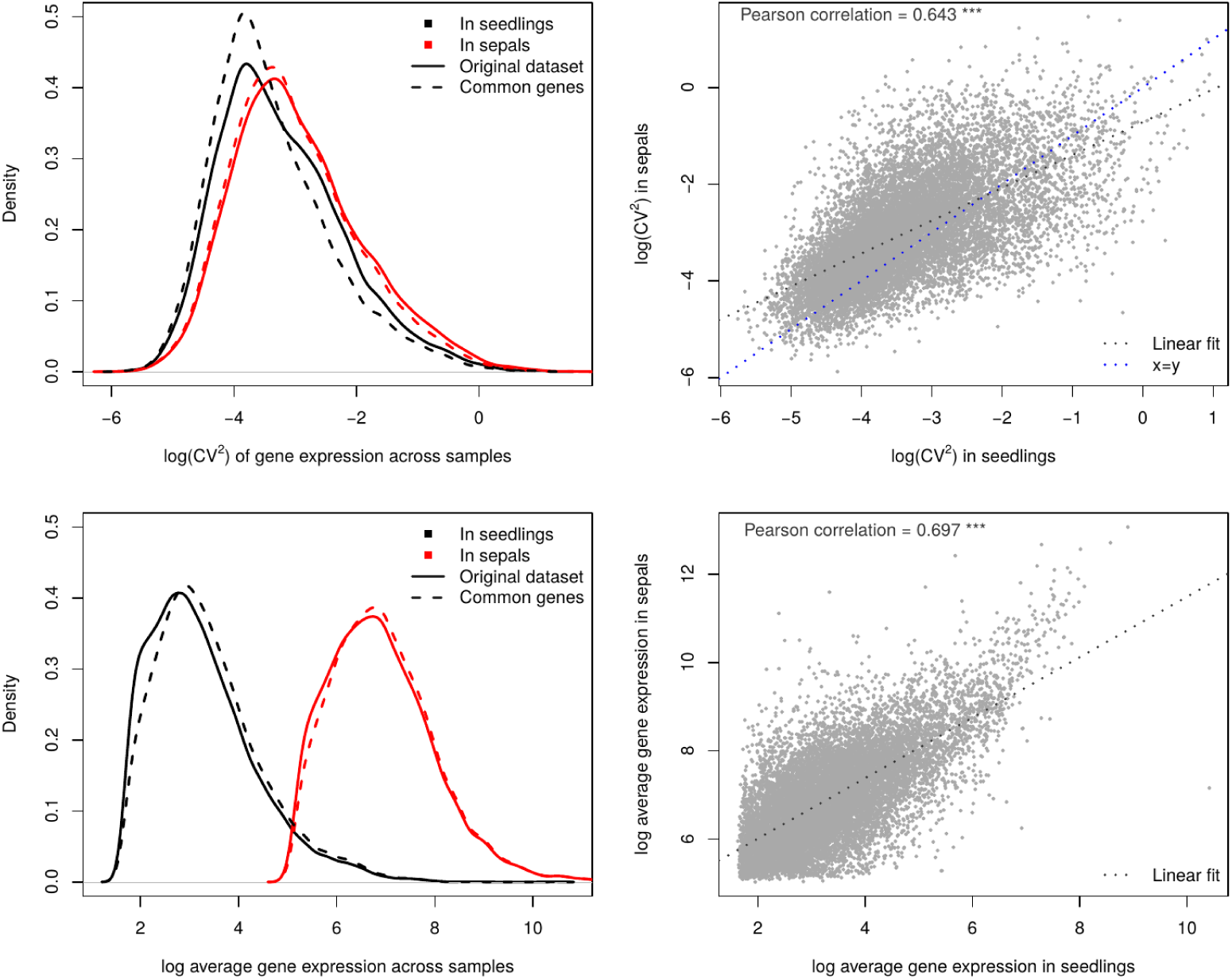
Validation of our RNA-Seq data by comparing squared coefficient of variation (top) and average gene expression (bottom) for our 14,085 genes (in red) with equivalent data (15,646 genes, in black) for Arabidopsis seedlings from Cortijo et al. (2019). Results are shown for each ”original dataset” (solid lines) and for genes in “common” (dashed lines) between both datasets (13,013 genes). Gene expression CV² values are comparable and have a strong correlation (top right). Mean gene expression values extracted from read counts show very similar distributions and a strong correlation (bottom right) but higher coverage in our case.

## Notes

### Competing Interest Statement

The authors have declared no competing interest.

### Summary of Updates

recommendation by PCI Genomics

